# Complete Three-Dimensional Spiral Ganglion Neuron Map in a Human Adult using a Deep Learning Pipeline

**DOI:** 10.64898/2026.11.10.765480

**Authors:** Christopher K. Giardina, Jennifer T. O’Malley, Abbie K. Hall, Anbuselvan Dharmarajan, Andreas H. Eckhard, M. Charles Liberman, Julie G. Arenberg, Alicia M. Quesnel

## Abstract

The quantity and distribution of spiral ganglion neurons (SGNs) greatly influence speech and hearing ability. Historically, approaches to quantify SGNs from archival human temporal bones (TBs) have utilized only every 10^th^ section to estimate the total SGN count. The current study evaluates a novel deep-learning pipeline for SGN quantification to produce the first complete three-dimensional SGN map of the human cochlea using every tissue section.

The reconstruction is based on one adult human cochlea in which all 421 consecutive, 20 um thick, celloidin-embedded sections were H&E stained, mounted and digitized. SGN nuclei were manually labeled in training sections which were subsequently used to create UNet++ models for SGN detection. These models were then applied to all other sections and compared to manual counts by three experienced histopathologists. After quantification, serial sections were aligned and SGNs were assigned percent distances along Rosenthal’s canal and characteristic frequencies using a modified Greenwood formula.

Models to automatically identify SGN nuclei from digitized tissue sections performed with great accuracy: SGNs were counted in 191 sections with 97% accuracy compared to manual counts. When tissue sections were systematically omitted, counting only every 10th section was determined sufficient to ensure a median inferred total count accuracy of 98%.

We present a new, flexible machine learning pipeline for identifying SGNs in digitized human TBs. Further, we show the first full 3D visualization of all 27,880 SGNs in an adult human cochlea, with associated tonotopic maps and basal-to-apical SGN distributions.

## INTRODUCTION

The quantity and distribution of spiral ganglion neurons (SGNs) have long been used as metrics in assessing the neural health of the human cochlea. Characterizing SGN survival from human temporal bones (TBs) has played a fundamental role in the understanding both of normal and abnormal changes observed in age-related hearing loss [1, 2] Ménière’s Disease, chronic otitis media, infectious labyrinthitis, ototoxicity [3], vestibular schwannomas [4] and outcomes analysis in cochlear implantation (CI) [5–12].

Until recently, SGNs have been manually quantified in human TBs by identifying SGN nuclei or nucleoli on Hematoxylin and Eosin (H&E) stained slides using a gridded eyepiece under a light microscope [13–15]. This process necessitates an expert histopathologist and patience, as standard 20-µm-thick modiolar slices can often contain hundreds of SGNs. In most archival human TBs in the United States, only every 10^th^ section is mounted and stained with H&E. The intervening sections are preserved for future stains, immunohistochemistry or molecular studies. For this reason, SGNs are counted on every 10^th^ section, and then numbers are scaled up by 10× and adjusted by a correction factor to infer a total count [15, 16]. If the goal were solely to infer a total count, modern approaches such as stereology [17] could provide excellent quantification much faster than manual counts, however this method does not retain any location data of SGNs, precluding analyses of density changes from base to apex. For this, a classic approach is to hand-count the SGNs through each cross section of Rosenthal’s canal (RC), mark these counts (per RC) on a two-dimensional (2D) spiral representation, and then sum the counts along four half-turn segments [2]. Using these methods, prior human TB studies have demonstrated that SGN counts are highest in infants or those within the first decade of life (35k-40k) [13] and steadily decrease with age at an approximate rate of ∼100 cells per year [2] or faster with concurrent inner ear disease [13]. Due to the tremendous effort in manual counting [18], there are no reports of SGNs on every serial section through a human TB in a manner which retains location and basal-apical orientation.

The recent digitization of histology slides for microscopy has created new opportunities in the analysis of TBs, in terms of automatic identification and three-dimensional (3D) reconstruction of SGNs. Wu et al [19] recently introduced a machine learning pipeline to identify SGNs that demonstrated high accuracy compared to manual counts (r²=0.945, F1 score=0.87±2) and assigned characteristic frequencies based on Greenwood’s function at the location on the organ of Corti spiral closest to the RC spiral, as computed in 3D space. Although the Wu model was accurate and effective, it required some fluency with Python and was a stand-alone application. We wanted to develop a flexible tool that could easily integrate SGN locations into commercial image-analysis software (such as Amira or Dragonfly). In this way, SGNs could be related in our ongoing studies of cochlear implantation to intracochlear pathology such as fibrosis and osteoneogenesis, to answer questions regarding the influences of CI insertion techniques [20], CI array type [21] and ultimately the effects of intracochlear pathology on speech ability and residual hearing preservation [22]. To apply this pipeline for either histopathologic or radiologic investigations, it needed to report SGN distributions either as a function of percent distance along RC or angular depth, respectively.

The current manuscript 1) introduces a new, flexible machine learning pipeline that can accurately quantify and map human SGNs using a coordinate system that can be co-registered with 3D maps of cochlear pathology, 2) presents the first complete three-dimensional SGN map of the human cochlea using every tissue section, and 3) validates a long-standing assumption that the total SGN count can accurately be estimated from only every 10^th^ tissue section.

## MATERIALS AND METHODS

### Archival TB Case and H&E Staining

An adult human TB was processed in the standard histologic fashion, except that *<u>all</u>* 421 consecutive, 20-um, celloidin-embedded sections were stained with H&E, mounted and digitized. This is in contrast to nearly all other archival TBs in the Mass Eye and Ear Otopathology Laboratory collection, in which only *every 10^th^* section is stained with H&E, mounted and digitized [15], creating a set of ∼40 total H&E stained sections.

The donor was a 71-year-old male with no known otologic disease in the study ear (left ear), and a history of Ménière’s disease in the contralateral ear (right ear). Audiometry seven months before death (Figure 1) revealed mild-to-moderate sensorineural hearing loss and excellent speech discrimination of the ear studied. The subject passed away at 71 years 10 months due to complications related to pneumonia and multisystem failure in the setting of decompensated hepatitis C. The post-mortem time to fixation was 14.5 hours, fixation occurred in 10% neutral-buffered formalin, and decalcification with EDTA (289 days) occurred before celloidin embedding and sectioning in the standard horizontal plane.

**Figure 1:**
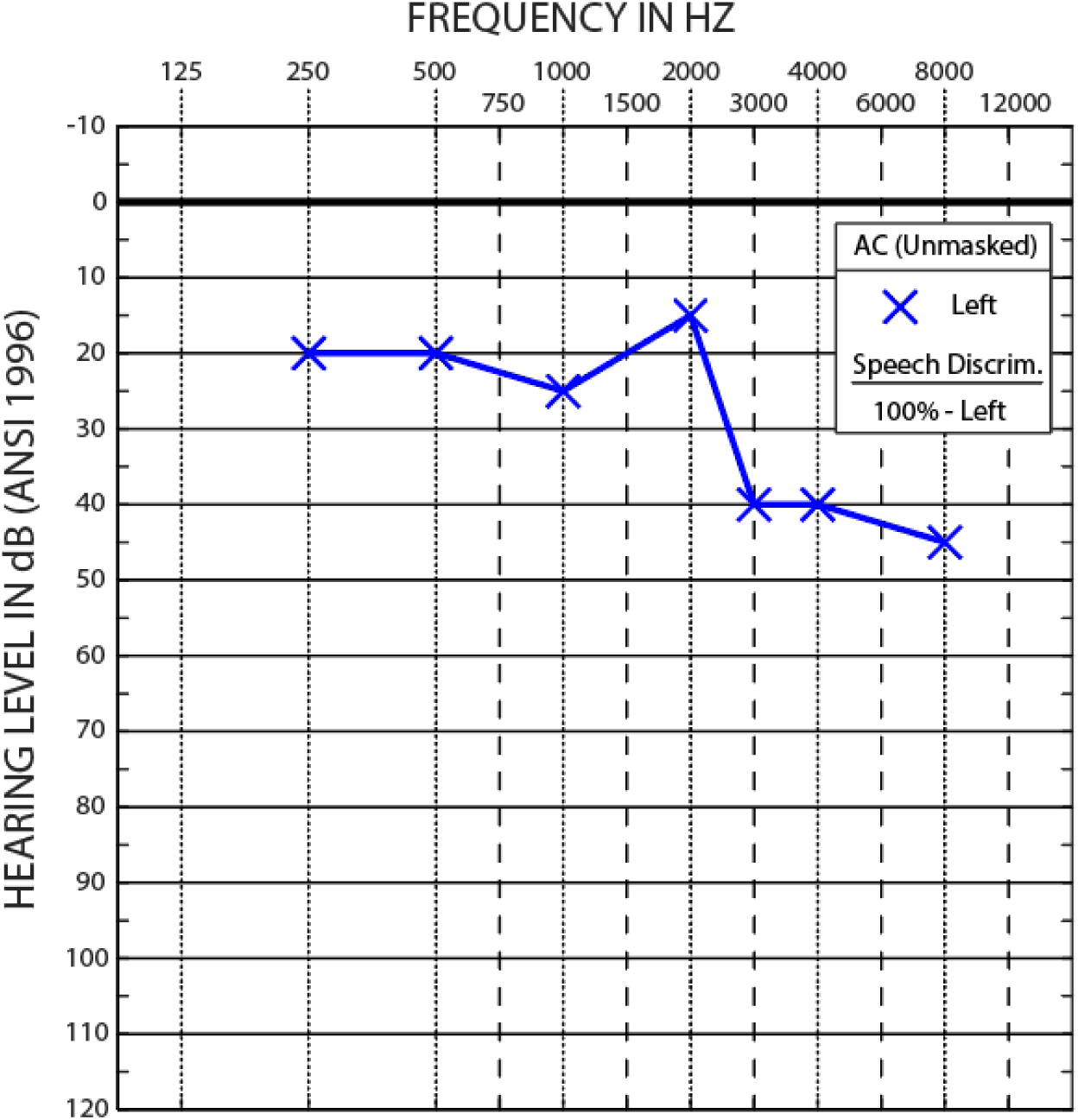
Audiometry and speech discrimination in the studied ear, 7 months prior to death at age 71. Hearing loss in the left ear was normal-to-mild below 3 kHz and mild-to-moderate above 3 kHz, with 100% speech discrimination assessed by Word Recognition Score (WRS). Contralateral ear (not shown) had a history Ménière’s disease, demonstrated severe to profound hearing loss, <50% WRS, and had Ménière’s profound pan-cochlear SGN loss.

### Slide Digitization with Single-Plane Leica AT2 Scanner

After staining, each section (421 total) was scanned into a digitized “eSlide” with a Leica Aperio AT2 scanner (Leica Microsystems, Wetzlar, Germany) fitted with a high-resolution 20× objective (NA 0.75). The scanner automatically acquires images at 4 separate focal levels, which it combines into a single continuous “eSlide”. While this automation simplifies many steps in digitization, it limits analysis to a single focal plane. Once fully scanned, the eSlide can be viewed at different magnifications, albeit at the cost of substantially increased file size, mimicking the user experience at the light microscope.

### 3D Reconstruction of Organ of Corti and Rosenthal’s Canal with Characteristic Frequency Assignment

As a first step, the 378 sections containing cochlear structures (of the total 421 sections scanned) were cropped at full resolution (8-bit RGB, 0.501-0.503 µm/pixel) using custom Matlab code to contain just the region of the cochlea. This stack of 387 TIFF images served as the starting point for alignment in 2D and subsequent reconstruction in 3D. Because loading nearly 400 high-resolution images requires ∼1 TB of RAM, a low-resolution stack (25x down sampling) was used for 2D alignment and then scaled up for subsequent analyses.

After alignment with a 2D affine transform using Amira3D (Thermo Fisher Scientific, Waltham, MA, USA), a custom Matlab script allowed a user to scroll through the aligned slices and mark the positions of the organ of Corti and RC (Figure 2A), similar to the method in [19]. For the organ of Corti path (“Hair Cell path”, HC), the intersection of inner and outer pillar cells (Figure 2B, black arrows) was tagged from base to apex, producing a 1,030-point spiral. For RC, the center of each bony canal (Figure 2B, blue arrows) was marked from base to apex, totaling 410 points. The 3D points along each spiral were smoothed with the *Smoothing Splines* function in Matlab [23] and vectorized into 1,000 equally-spaced steps. For each step along the HC path, an associated Characteristic Frequency (CF) was assigned using a modified Greenwood equation (1) [24]. Similarly, for each step along the RC path, an associated CF was assigned using a best-fit formula derived from 3D synchrotron imaging [25].

**Figure 2:**
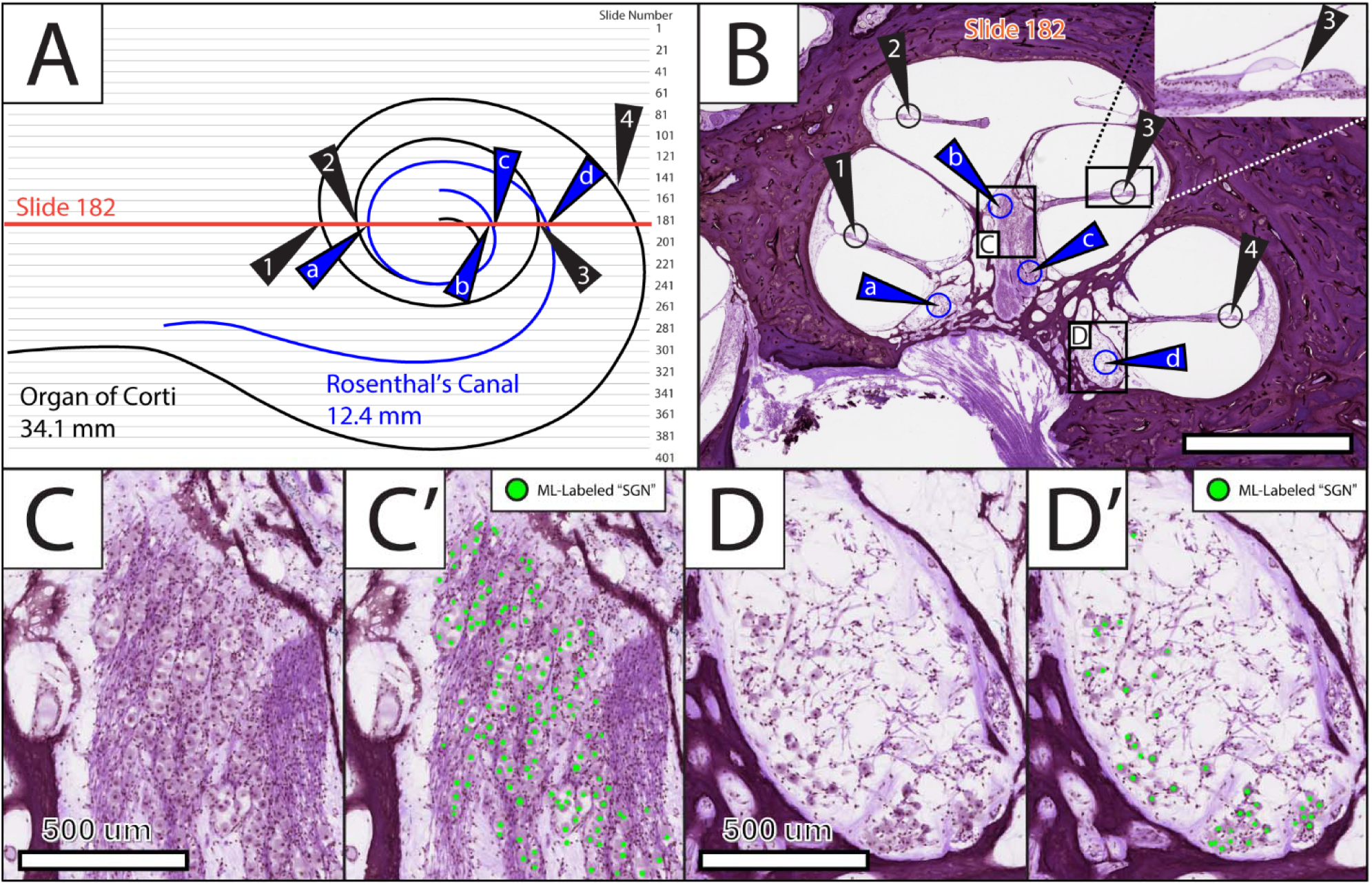
Annotating Organ of Corti and Rosenthal’s Canal for 3D reconstruction and ML pipeline. **(A)** 2D spiral representation of Rosenthal’s Canal (RC) and the Organ of Corti path for all histology slides in the stack, with Slide 182 labeled. **(B)** On Slide 182, Organ of Corti landmarks 1-4 and RC landmarks a-d can be seen (bar = 2 mm). Raw slide and machine learning (ML) labeled regions of high SGN density **(C,C’)** and low SGN density **(D,D’)** regions of RC.

Equation 1: Assigning CF by Percent Distance along Organ of Corti [24]

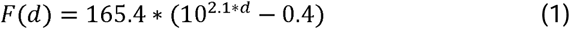

Equation 2: Assigning CF by Percent Distance along Rosenthal’s Canal [25]

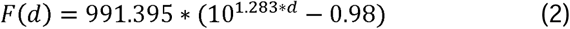

The HC and RC paths were then upscaled 25 to match the full-resolution images, allowing for full 3D visualization of all landmarks in Matlab. This shared coordinate registration allows for integration of 3D intracochlear anatomy plotted or mapped in other software packages to be plotted alongside the HC path, RC path, and SGN points from the core workflow.

### Supervised Deep-Learning Segmentation Model for SGN Identification

The identification of SGN nuclei was facilitated by training and applying a custom deep-learning segmentation model within Dragonfly 3D World 2024.1 (Comet Software, Montreal, QC, CA). The overall workflow involves 1) importing single-channel (red, green, or blue) images into Dragonfly, 2) hand-annotating pixels of SGN nuclei against background into a binary “mask” for training, 3) training the deep-learning model to identify SGN nuclei, 4) validating the model against hand-labeled slides not used in training, and 5) applying the model to all other slides in the case.

First, uncompressed TIFF images of each cochlear section were imported into Dragonfly as three discrete color channels, of which only green was used for training, as this channel had the best contrast for the nucleus in comparison to other structures.

For model training, only a single mid-modiolar slice was hand-annotated with SGN nuclei, but later models utilizing three or even eight annotated slices were created with no appreciable difference in accuracy. Parameter optimization using the Deep Learning Tool within Dragonfly allowed for streamlining of both training data augmentation as well as parameter fitting (specific variable settings in Supplemental Table 1). While a standard UNet encoder/decoder architecture [26] would likely have been sufficient, we opted to implement a UNET++ (“Nested UNet”) architecture [27], which takes advantage of dense [28], nested skip pathways between encoder and decoder feature maps. Once trained, the model was applied to all other images in the stack, selectively around areas where RC was tagged (Figure 2C,C’ and Figure 2D,D’), generating a binary pixel mask for each class (again 0=background, 1=label). The Connected Components function within Dragonfly combined contiguous pixel groups into a list containing centroid X coordinate, centroid Y coordinate, and voxel surface area of each pixel cluster.

### Confocal Imaging

To assess the extent to which the eSlide counts capture all the SGNs within a tissue section, four consecutive microscope slides with H&E stained sections from the same TB case were imaged using a Leica TCS SP8 confocal (Leica Microsystems, Wetzlar, Germany) equipped with a 20x glycerol immersion lens (NA 0.75). Unlike the eSlides—which contain a compressed single focal (z) plane and therefore do not fully resolve the entire thickness of the section, posing a risk of missing out-of-focus nuclei—confocal z-stacks were acquired at 1.5 µm intervals. Eosin fluorescence was excited at 561 nm and emission was detected between 564 and 668 nm with a photomultiplier tube (PMT). All images were acquired at 4096 × 4096 pixels. Imaging parameters were kept constant for all samples.

### Manual SGN Counts

To assess the accuracy of machine learning (ML) derived SGN counts, independent counts from three experienced histopathologists were used at each stage. For the ML-training eSlide and the ML-validation eSlides (every 10^th^), counts were made using custom Matlab code which logged coordinates, which could then be compared among the raters. Additionally, for the ML-validation slides, custom Matlab code allowed categorization of ML-tagged nuclei as ‘true positive’ or ‘false positive’, while logging nuclear areas (in pixels) to determine a cutoff for ML-segmented noise. For the four consecutive slices which had confocal imaging (Figure 3C’-F’), SGN nuclei counts on eSlides were compared to counts from the confocal z-stacks after segmenting nuclei with Amira 3D.

**Figure 3:**
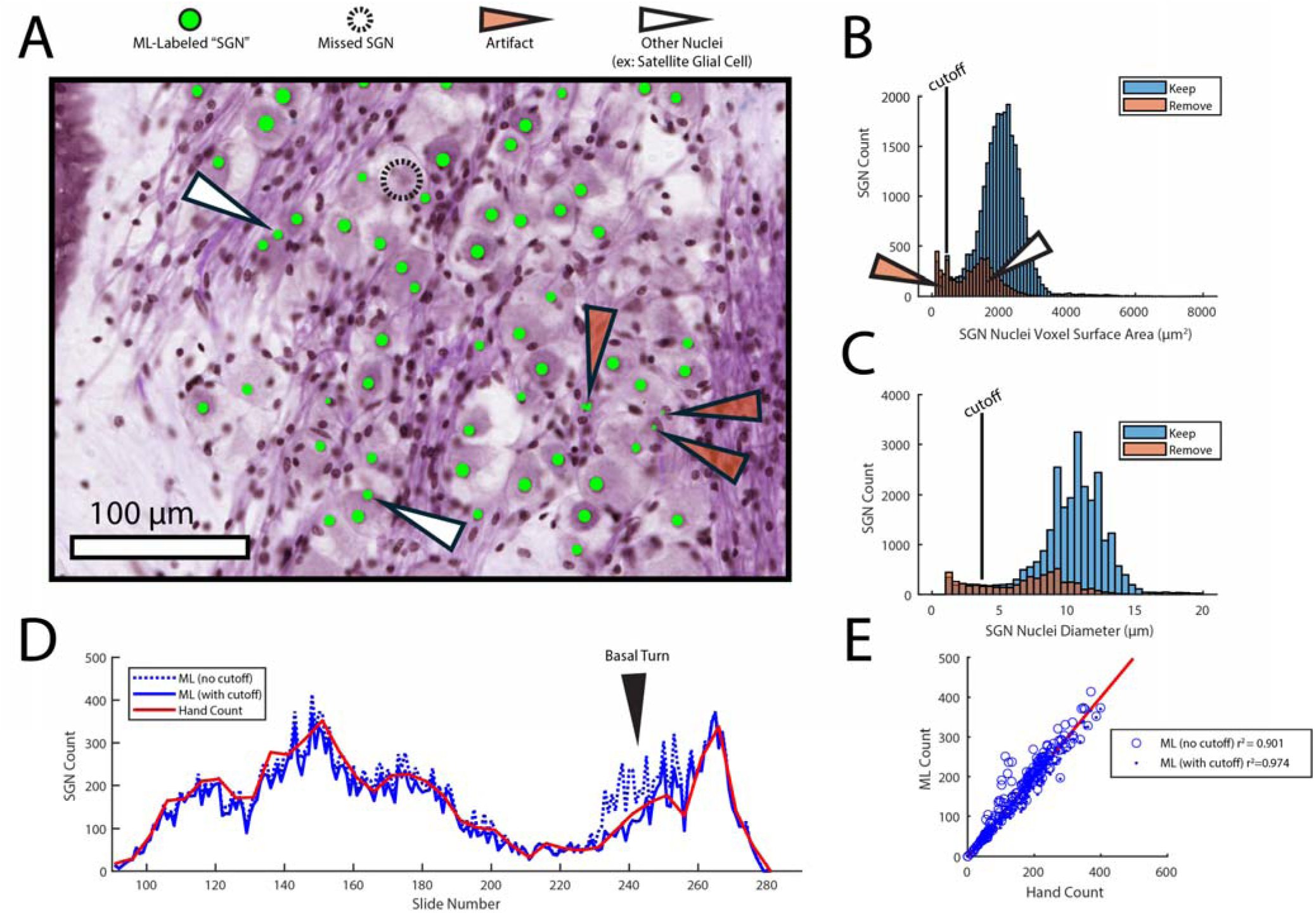
Accuracy of Segmentation Model for SGN Detection. **(A)** High magnification region of eSlide with raw SGN masks applied by ML (green). A false negative (dashed circle) appears to be slightly out of focus. Two types of fal e positive – noise artifact (orange arrow) vs. Schwann cell nucleus (white arrow) - are noted. **(B)** Histogram of ML-derived SGN segmentation areas, with observer-derived false positives indicated in orange and a size cutoff of 407 µm ^2^. **(C)** Same data as B, except plotting diameters instead of areas. **(D)** Comparison of manual vs ML-derived SGN counts as a function of slide number (See Fig. 2A). Black arrow indicates slide range containing the basal turn (refer to Fig. 2A for cochlear orientation by slide number). **(E)** Accuracy of SGN counting improved from 90.1% to 97.4% when using cutoff.

### Assigning SGNs to an Angular Insertion Depth

Once SGNs were labeled on each image, a 3D cochlear reconstruction with HC, RC, and SGNs was created. At this stage, each SGN was assigned a CF by finding the segment of the HC path closest to the RC and assigning the CF of that region [19]. As a final step in 3D registration, a modiolar axis was identified in order to determine the angular coordinate of each SGN. A custom Matlab script allowed for rotation and translation of all points (HC, RC, SGN) until the cochlea was oriented with the modiolus orthogonal to the projection plane, as seen in Figure 2a, and similar to the ‘Cochlear View’ seen with radiology imaging [29, 30]. With this orientation, the function *arctan2d()* in Matlab calculates the angle of each of the 1000 segments along the HC and RC paths. Additional code was necessary to calculate positions in the “hook” region of the spiral. SGNs were assigned an angle based on the angle of the closest RC segment. Histograms of SGN counts by angle were generated in Matlab with the *histogram()* function using angular bin sizes of 5, 10, 15, 30, 45, 60, and 90 degrees.

### Artificial Removal of Slices

With all SGNs in the cochlea assigned both a CF and cochlear angle, the next step was to artificially remove SGNs on predefined slides to emulate situations where, for example, only every 10^th^ slide had been stained and counted. This situation poses two related questions: 1) what is the effect of slide spacing on accuracy, and 2) does the specific starting slice number matter. For a slice spacing of 10, for example, there are 10 groups of slides which could be quantified-each with a different starting slice number (ex: 1,11,21,31 vs 2,22,32,42, and so on). Total accuracy (total count) was assessed by comparing the estimated count of each separate group (multiplied by 10) to the true known total. This was repeated for all additional slice spacings of 2, 3, 5, (10), 20, and 40. To incorporate the analysis of both slice spacing and starting number on angular bin counts, a similar analysis was performed for each angular bin rather than a total. In this way, it would be possible to determine if errors from slice spacing and bin size was disproportionate in any angular region of the cochlea. Median errors and maximal errors per bin were evaluated for each combination of slice spacing (2, 3, 5, 10, 20 and 40) and angular bin size (5, 10, 15, 30, 45, 60, and 90 degrees).

## RESULTS

### Accuracy of Segmentation Models to identify SGNs on individual eSlides

The core technique used in this study was the automated identification of SGNs on eSlides using machine learning. The training eSlide used to create the model incorporated regions of both high and low cellularity, which allowed the UNet++ to identify SGN nuclei on new slides with both high (Figure 2C) and low (Figure 2D) density. To evaluate accuracy and characterize errors, visual inspection of raw output and a comparison of ML counts to hand-counts on every 10^th^ slide were made (Figure 2). The raw output of the segmentation model is a binary pixel-by-pixel classification. When clusters of labeled pixels are adjacent to each other, the ‘connected components’ function combines this pixel region into a single ‘SGN nucleus’ label, with both location (centroid X and centroid Y) and surface area (in voxels) data. Visual inspection of a typical ML-labeled area (Figure 3A) reveals the typical raw output from the applied segmentation model.

Green areas are often correct (per visual inspection), however examples of false positives and false negatives can be seen. A nucleus was missed during ML labeling (Figure 3A, dashed ring), however this indistinct element may also have been missed by expert histopathologists. Additionally, two types of false positives could be seen. Some small ML-labeled puncta (Figure 3A, orange arrowheads), are pixel regions associated with acellular dark patches, and others (Figure 3A, white arrowhead) are nuclei of non-neuronal cells, i.e. the satellite cells ensheathing the SGN cell bodies or the Schwann cells around the axons. Of note, satellite and Schwann cell nuclei are smaller than SGN nuclei; therefore, we considered a pixel-area cutoff to exclude the smaller nuclei.

To determine whether a lower pixel-area cutoff could be used to remove ‘noise’ and also determine the extent of overcounting of nuclei from non-SGNs, every ML-labeled SGN on every 10^th^ slide was manually reviewed. As expected, the voxel areas of the SGNs identified as false positives by the observers showed a bimodal distribution (Figure 3B), which was still present when calculating for fitted nuclear diameter (Figure 3C). The ‘diameter’ in Figure 3C is that of the ML-classified region and does not necessarily match the full diameter measured under light microscopy. The raw count across all sections (Figure 3D, dashed line) is higher than the hand-count (red line), however when introducing a lower area cutoff of 407 µm^2^ to reduce the noise (orange labels), the correlation between raw ML count and manual count increases from r^2^ of 0.901 to 0.974 (Figure 3E).

### Correction Factor is not necessary when counting on eSlides

An essential consideration when counting cells across tissue sections is correcting for double counting when cells are bisected during sectioning. For this reason, SGN counts are typically scaled down by the Abercrombie correction factor [31] computed based on the ratio of the size of the counted element (e.g. nucleus or nucleolus) to the thickness of the section [15]. Recent SGN studies have analyzed correction factors for counts made under light microscopy [32], however no formal study has been made for eSlides, where SGN nucleoli are difficult to see, thus counts are made on nuclei only. When four consecutive sections through a cluster of SGNs in the upper basal turn (Figure 4A) were imaged as a confocal series with 1.5 micron spacing in the focal plane (Figure 4B), a total of 25 SGNs was observed, of which 5 nuclei were split across sections. When the four sections were each counted under a light microscope, taking care to focus through the entire thickness, 29 nuclei were observed. Thus, the empirically derived correction factor for light microscopy is 0.86, i.e. similar to the 0.66 value derived from the Abercrombie formula. When utilizing eSlides, viewing only a single focal plane presumably near the middle of the section thickness, no nuclei span two adjacent eSlides (Figure 4C-4F). Thus, when the four sections were counted using eSlides, 25 nuclei were observed, exactly as seen from through-focus confocal analysis.

**Figure 4:**
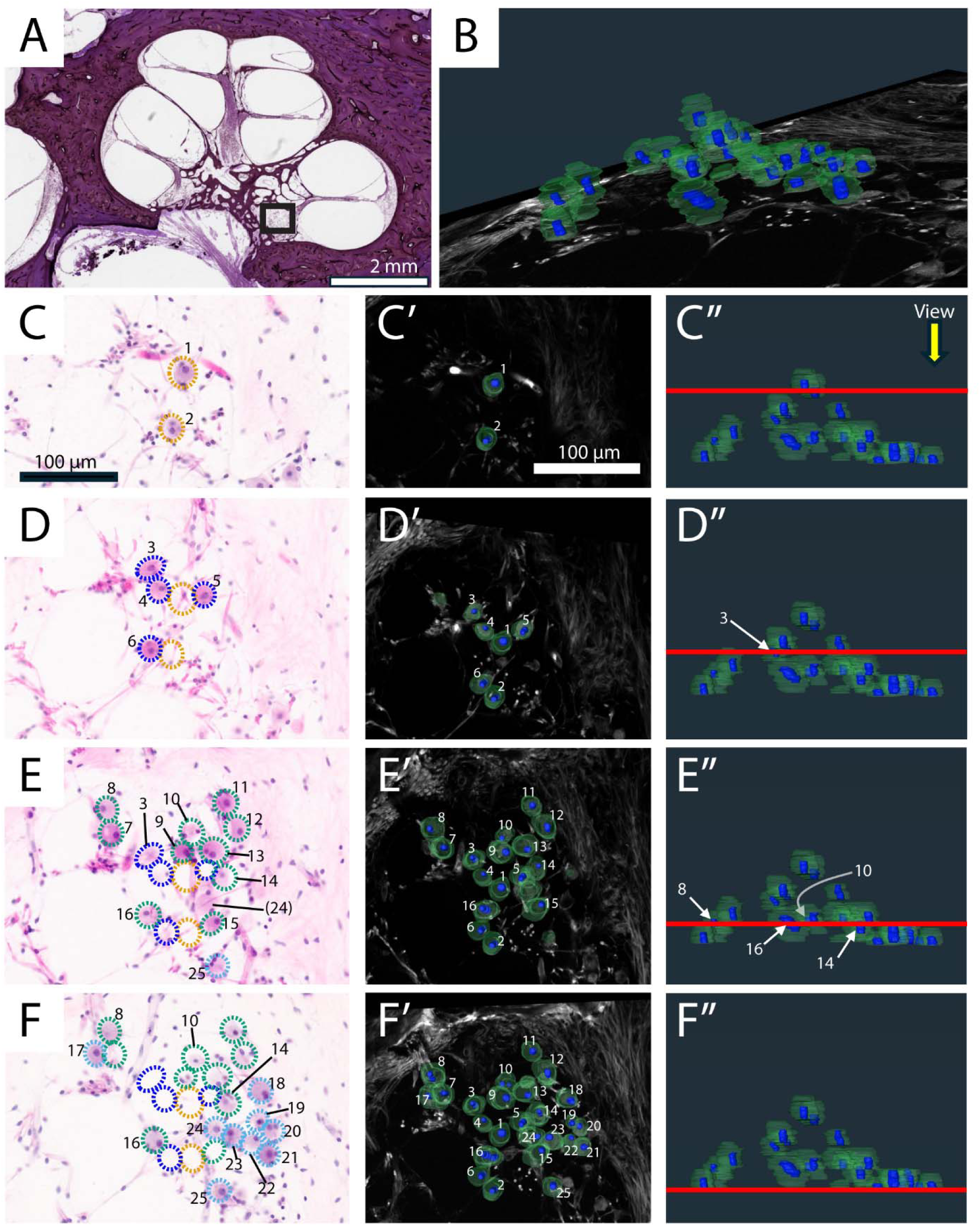
Evaluation of SGN nuclei across 4 consecutive slices when using eSlides or confocal imaging. **(A)** First eSlide and region of Rosenthal’s Canal (box) for analysis. **(B)** Volumetric rendering of SGN cell bodies and nuclei from 60 sequential focal planes across 4 slides produced by confocal imaging. **(C,D,E,F, Left column)** 4 sequential eSlides zoomed into highest resolution, with SGNs numbered. **(C,D,E,F, Center column, Right column)** Volumetric renderings at top-down views of different Z depths (red line), with same SGNs numbered as Left column.

### SGN Quantity and Distribution in an Adult Human

Without a correction factor, the raw count of SGN nuclei in this TB was 27,880. After a full 3D reconstruction (Figure 5A,5B), the overall length of the ‘HC’ Organ of Corti path (Figure 6A, gray line) was 34.1 mm and spanned −7 to 825 degrees (∼2.35 turns) around a RW reference angle of zero degrees. The ‘RC’ path (Figure 6A, red line) was 12.7 mm, and spanned −5 to 780 degrees about the modiolar axis. When quantifying SGNs in angular bins of 30 degrees (Figure 6B), SGN counts were highest at the base and generally decreased towards the apex. However, because the angular lengths of RC are greater at the base than apex (black line in 6B, and alone in Figure 6D), SGN density was normalized to this RC arclength (Figure 6E) and revealed a more uniform distribution from base to apex with lower densities at either end. Several CF values, as inferred from the distance along RC (Equation 2), are shown in angular bins as a reference. To compare the nuclei counts in this TB with other studies, SGN counts are also shown as a function of RC distance (Figure 6C) and per RC Segment (Figure 6F, where Segment I: up to 90 degrees, II: 90-270 degrees, III: 270-450 degrees, IV: greater than 450 degrees).

**Figure 5:**
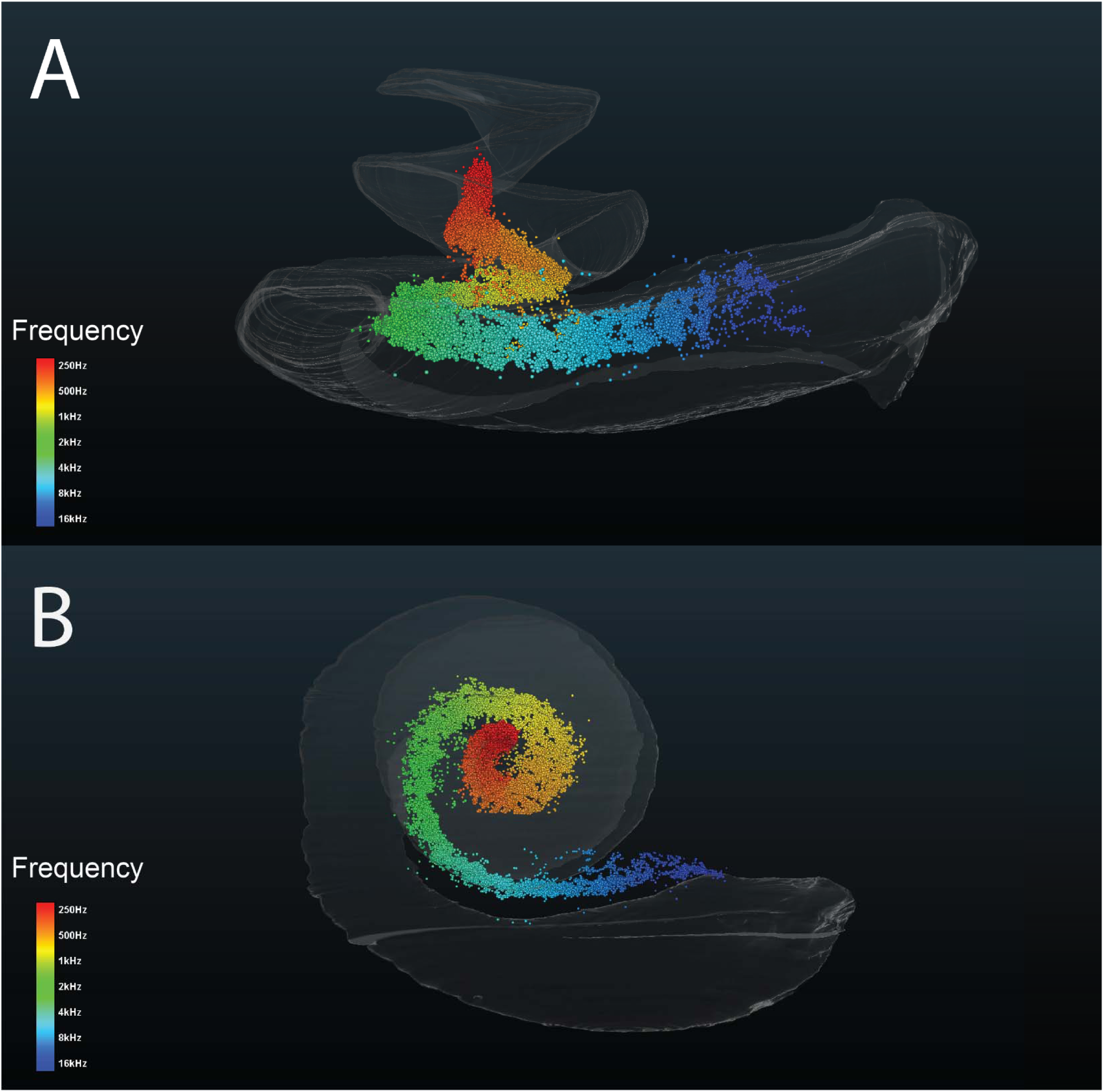
Three-dimensional map of all 27,880 SGNs in the cochlea, with assigned CFs. **(A)** Lateral view and **(B)** top-down view of all 27,880 SGNs with assigned CFs indicated by color legend. Gray volume is Scala Tympani.

**Figure 6:**
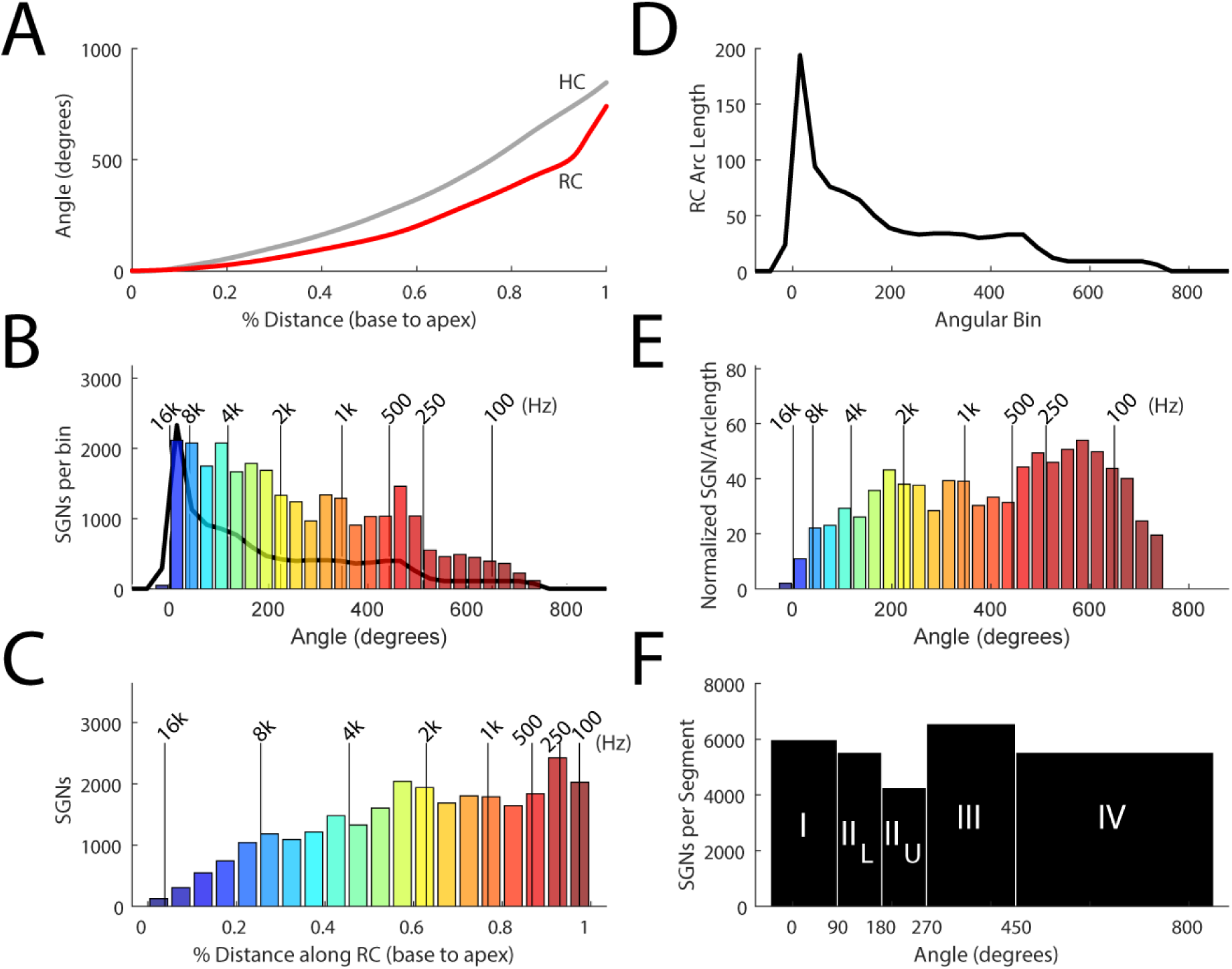
Distribution of all 27,880 SGNs. **(A)** The Hair Cell (HC) path (along the Organ of Corti) was 34.1 mm and spanned an angular range of –7 degrees to 825 degrees (relative to the round window), whereas the Rosenthal’s Canal path (RC) was 12.7 mm and spanned an angular range of –5 degrees to 780 degrees. **(B)** SGN counts per 30 degree angular bin about the modiolar axis, colored by CF, with black line showing the RC arclength of each 30 degree angular bin **(line alone in D)**. **(C)** SGN counts per 5% distance segment along RC. Since angular areas account for a much larger linear length in the base and smaller linear length in the apex compared to % linear distance, SGNs counts appear very different when plotted by angle (B) vs. distance(C). One way to account for this is to normalize angle/ length, as shown in E. **(E)** SGN count per angular bin, normalized by RC length. **(F)** SGN count per half-turn segment. kHz=kilohertz

### Slice Spacing and Angular Bin Size to Accurately Approximate Quantity and Distribution

The process of quantifying every section allowed for the relationship between slice spacing and accuracy to be characterized by removing slices, i.e, leaving every 3^rd^, 5^th^, 10^th^, 20^th^ or 40^th^ slice (Figure 7). The first groups (starting slice number=1) for several key slice spacings are depicted in Figures 7B-F, demonstrating cross sections of the equally-spaced remaining slides. Counts per artificially induced spacing were scaled and compared, to determine accuracy (Figure 7G). When the spacing increased, the number of possible starting positions also increased (i.e.: spacing of 2 only has 2 possible sets of slides, spacing of 10 has 10 sets). When investigating all combinations of slice spacing and starting section number, errors in estimated total count greater than 10% were not observed in more than half of the groups until slice spacing of 20 or greater was used (Figure 7G).

**Figure 7:**
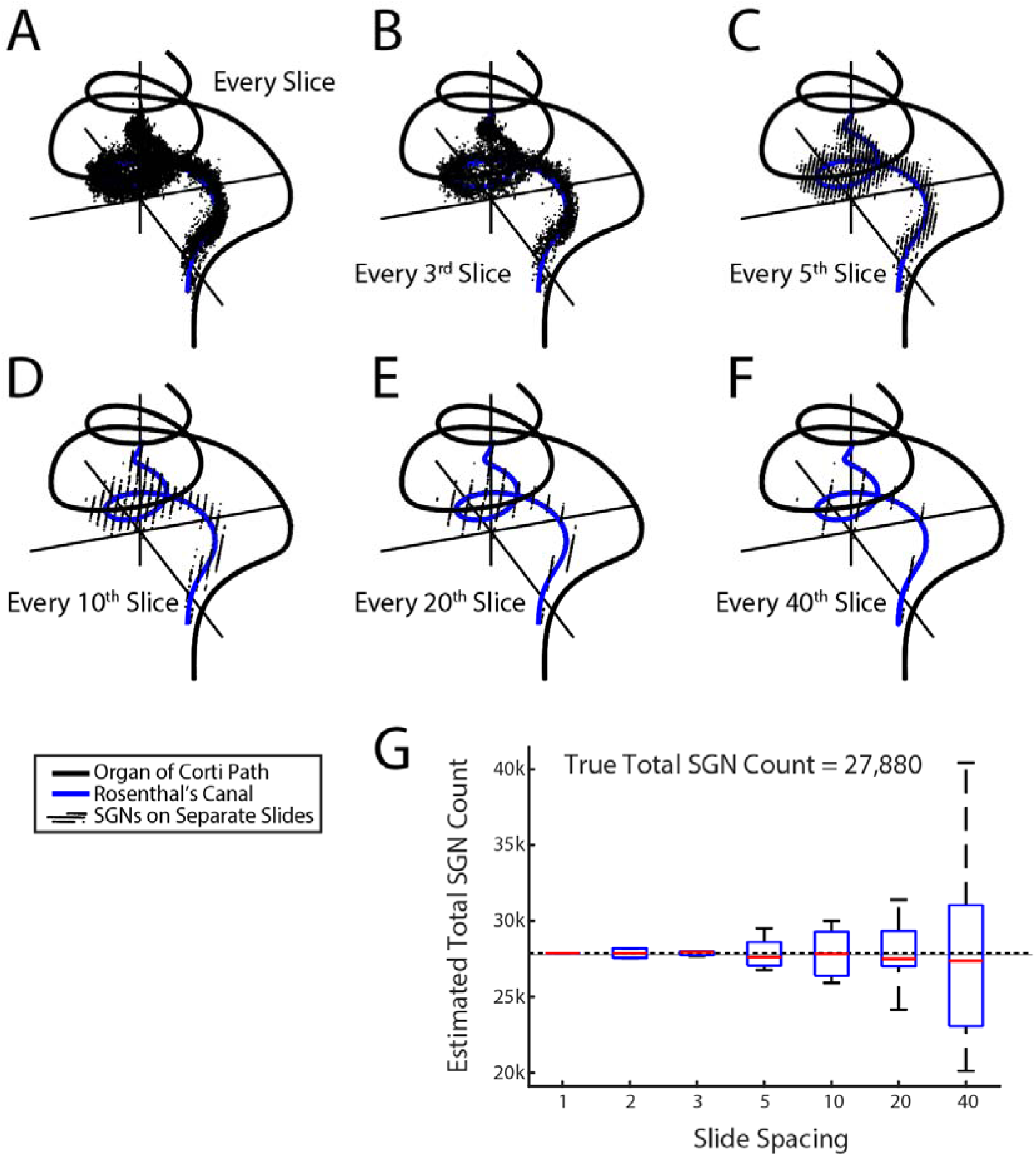
Effect of slice spacing on accuracy of total SGN count. **(A)** Cochlea with SGNs from every slice. **(B)** SGNs from every 3rd slice when starting on slide 1. **(C)** SGNs from every 5^th^ slice when starting on slide 1. **(D)** SGNs from every 10^th^ slice when starting on slide 1. **(E)** SGNs from every 20^th^ slice when starting on slide 1. **(F)** SGNs from every 40^th^ slice when starting on slide 1. **(G)** Accuracy and variance when estimating total SGN count from all slice spacings and any starting slide number.

In addition to accuracy in total count (Figure 7), we explored the effect of section spacing (and starting slice) on accuracy when quantifying angular bins (Figure 8A-C), to determine if any regions were disproportionately affected by slice under-sampling, and, if so, was there an ‘ideal’ angular bin size to minimize these differences. The median error of all angular bins (blue dashed line on right of 8B) increased with slice spacing but was below 100 SGNs when slice spacing was 10 or lower. Maximal error was usually in the basal turn around 45-90 degrees (black lines on right of 8B). As shown in Figure 8C, median bin error increased when either slice spacing increased (Figure 8C, different colors) or bin size increased (horizontal axis), and smaller bins generally afforded closer estimates to the true counts. An evaluation of median error per distance bin (rather than angular bin) was also performed (Figure 8D), revealing similar variances as those in 30-degree angles (Figure 8B), however there was less variance at the base and larger variance at the apex.

**Figure 8:**
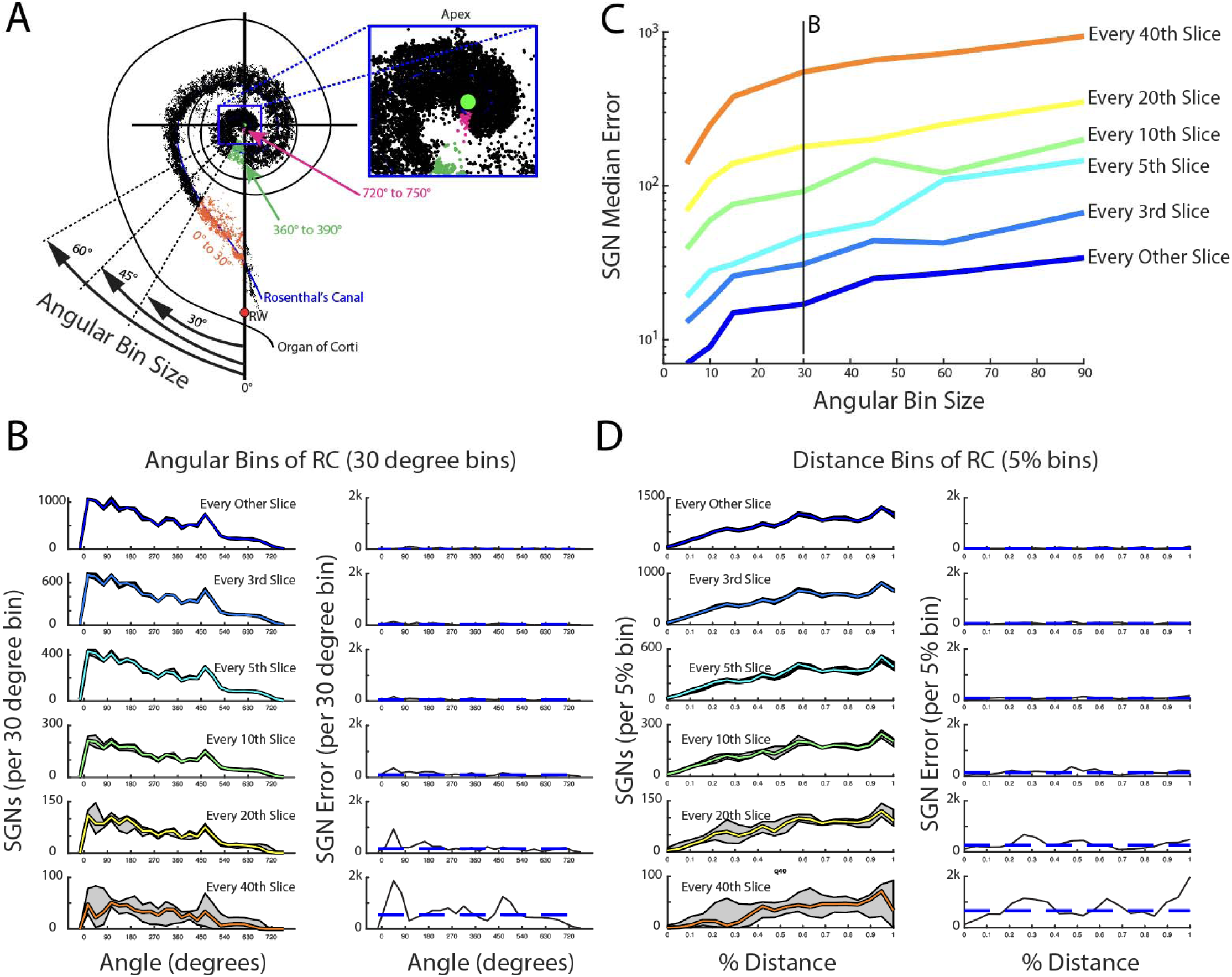
Effect of slice spacing and angular bin size on accuracy of angular SGN counts. **(A)** Modiolar vie of cochlea demonstrating angular assignment of SGNs based on angular bin sizes of either 30 degrees of 60 degrees. **(B)** When selecting angular bins of 30 degrees, variance in accuracy of each angular bin (gray areas) and median SGN error of all bins (blue dashed lines) increase with slice spacing. **(C)** Systematic analysis of angular bin size and slice spacing on median errors in estimation. **(D)** SGN error as a distance function along RC rather than angle.

## DISCUSSION

Automated techniques to quantify SGNs from eSlides were comparable in accuracy to the gold-standard of manual counting on eSlides, while adding significant benefits associated with digital volumetric reconstructions. Here, the overall processing pipeline allowed for identification, localization along RC, modiolar angle assessment, and CF assignment of all 27,880 SGNs by selecting cochlear landmarks on each eSlide.

### Identifying SGN Nuclei on individual eSlides

Identifying SGN nuclei from sections of this particular human TB was both accurate and efficient when using segmentation modeling approaches to digitized H&E-stained tissue sections. While other quantitative otopathology studies in animal models have also used computer vision approaches to localize HCs [33] or SGNs [34] with great accuracy, these studies used fluorescently tagged cells (affording higher contrast than H&E) and had short post-mortem times (minimizing cellular degradation). In human TBs, H&E staining is not uniform between cases and longer post-mortem times in some TBs may lead to SGN necrosis. Despite the wide range in SGN staining intensity between cases, SGNs within a given TB tend to have a similar appearance. For this reason, our approach of annotating a single slice to train the ML model should be successful in other cases.

While the raw ML output often included small puncta that were not nuclei (Figure 3), for larger puncta the errors made by the segmentation model were not substantially different from those made by observers. For example, nuclei from satellite cells were sometimes identified by the model (Figure 3A). As two satellite cells typically ensheathe the soma of each SGN, reducing the eSlide to a single focal plane could sometimes lead to false positives. A previous ML-driven SGN segmentation also found satellite cells were sometimes difficult to differentiate from SGNs on eSlides, though overall accuracy was not substantially affected [19]. Other differences when identifying SGNs on eSlides rather than under light microscopy arise from the loss of morphological detail when imaging only a single focal plane. This makes identifying SGN nucleoli and peripheral/central axons on eSlides unfeasible. However, hand-counts of HCs or axons under light microscopy can be integrated into a 2D model and matched by CF, an approach which showed the fraction of SGNs surviving without a peripheral axon increases with age, reaching more than 50% in the apex and 66% in the base [19].

### Correction factor for SGN nuclei on consecutive eSlides

When tissue is serially sectioned, cell counts across slices can exaggerate the true count because some cells are split and thus double-counted. Abercrombie [31] defined a simple formula to compensate for this overcounting, based on the ratio between the size of the counted element (e.g. cell nucleus or nucleolus) and the section thickness. Historically, a correction factor of 0.9 was adopted by most groups quantifying SGNs [13, 16, 35], because they were only counting SGNs with a nucleolus (diameter = 2.5 μm). If counting nuclei, with a mean diameter of 10.6 μm and a slice thickness of 20 μm, the correction factor should be ∼ 0.65, prompting later studies which counted nuclei to use a range of smaller correction factors [1, 3, 6, 36]. A recent empirical derivation of SGN correction factors [32] using digital reconstructions similar to the approach used here, determined the correction factor for nuclei was 0.91. One key criterion that is not always enunciated in SGN studies is whether cells are counted when only a sliver of the nucleus is visible. If only full-size nuclei are counted, the Abercrombie correction will not be appropriate.

In the current manuscript, a correction factor was empirically derived to characterize double-counting on eSlides rather than under light microscopy (Fig 4). In addition to double-counting, the effective down-sampling of the slice thickness into a single focal plane introduces the possibility of undercounting. In a set of 4 sequential slices, imaging of 80 focal planes revealed 5/25 SGN with nuclei which spanned across adjacent slides (Figure 4C’’-F’’). While this expected overcounting is close to that predicted by the Abercrombie formula, our localized analysis showed no double counting or undercounting in the ML-driven identification (Figure 4C-F). It appears that the nuclei near the section boundaries must be out of focus enough that they are not identified by the model. Thus, we conclude that a correction factor should not be applied when counting SGN nuclei with our methodology from digitized slides. The single plane of focus of the digitized slides (in comparison to the ability to scroll through the depth of the slide using standard microscopy) and application of this machine learning program to 20 μm slides effectively negate the issue of double counting nuclei. It is possible that findings may differ in areas of increased neuronal density, however, such as in cochleae from pediatric donors or younger adults, and that a correction factor may be applicable in those cases.

### Total SGN count

The total SGN count of 27,880 in an adult with a normal age-adjusted audiogram is consistent with several prior reports of normal-aging adults based on analysis of every 10^th^ section, where counts range from 23,000 to 39,000 [37] and 20,000 to 37,000 [13]. Our count is very similar to the mean value of 25,795 predicted for a 71 year old from a study of 100 age-graded temporal bones [2], while our total count was slightly lower than the 29,000 to 40,0000 range found in another study [38], their approach counted nucleoli and utilized a different correction factor, making direct comparison less straightforward. When evaluating the SGNs per RC segment (Figure 6F), counts were slightly higher in the base than in a prior study, however not substantially different in more apical regions [2]. Given that the case we studied had a 100% score on word recognition, it is unsurprising that slightly higher counts were observed.

### Angular SGN counts and implications for the study of TBs with Cochlear Implants

Because SGNs are the primary target for stimulation by cochlear implants, a reasonable hypothesis is that SGN survival should influence speech performance. While one within-subjects study found that higher SGN counts were associated with better CI performance in bilaterally implanted subjects [6], most CI studies have found no correlation between total SGN count and speech performance [39, 40] and some studies have even reported a negative correlation [7, 8]. These investigations have utilized total SGN counts or SGN per quadrant/segment of RC and have not taken into account the location of electrodes relative to surviving SGNs. A more fine-grained analysis of these adjacencies is now possible with the type of 3D reconstructions created in the present study.

Based on the data presented here, studies of SGN distribution from every 10^th^ section can estimate SGN counts with minimal error. We found that every 10^th^ section was sufficient to ensure an SGN median error below ∼100 SGNs per 30-degree bin. Importantly, this error was highest within the first 90 degrees, corresponding to the same basal region that may overlap with areas of surgical drill-out [21] or fibrous formation [22]. SGN errors in the basal turn may be minimal if the true counts here are also low.

Histological features seen in CI cases or other pathologies will need to be incorporated alongside SGNs in order to more fully understand the relationship between cochlear histopathology and audiometric performance. As human TB collections from donors continue to grow, digital solutions with novel approaches continue to advance our understanding of otopathology, disease, and how cochlear implants work.

## CONCLUSION

We present a flexible, machine learning pipeline to identify SGNs from digitized H&E-stained tissue sections, which was developed and validated based on comparison to the current gold-standard of SGN identification by expert histopathologists with consensus review. This pipeline identifies the number and three-dimensional distribution of SGNs and reports this distribution in either angular insertion depth or percent distance along Rosenthal’s canal. Utilizing a specimen in which we H&E stained and digitized every 20 μm thick section through the cochlea and applying this new, validated machine learning pipeline, we were able to produce the first complete human SGN map in a single cochlea showing the three-dimensional distribution of all 27,880 neurons.

## Acknowledgments

The authors are appreciative of the donors who have donated their temporal bones and the families who assisted with donations through the NIDCD National Temporal Bone Registry. We thank MengYu Zhu, Diane Jones, and Barbara Burgess for the processing and digitization of histological specimens.

## Author Contributions

CKG, MCL, JGA, and AMQ conceptualized and designed the study. CKG, JTO, AD, AHE, and AMQ contributed to methodology and validation. CKG performed the analyses and drafted the manuscript. CKG and AKH created the figures. CKG, AKH, JTO, AHE, MCL, JGA, and AMQ reviewed and edited the manuscript. JGA and AMQ supervised the project. All authors reviewed and approved the final manuscript.

## Data Availability

The data that support the findings of this study are available from the corresponding author upon request.

## Code availability

The code used in this study are available upon request.

## Artificial Intelligence

A Unet++ model trained and implemented in Dragonfly 3D World (version 2024.1) was used to quantify spiral ganglion neurons in this study. No generative artificial intelligence was used for manuscript preparation.

## Ethics Approval

Consent for use of the temporal bone in this study was provided by the donor. This study was approved by the Institutional Review Board of Mass Eye and Ear (IRB#2021P001593).

---

**Supplemental Table 1:**
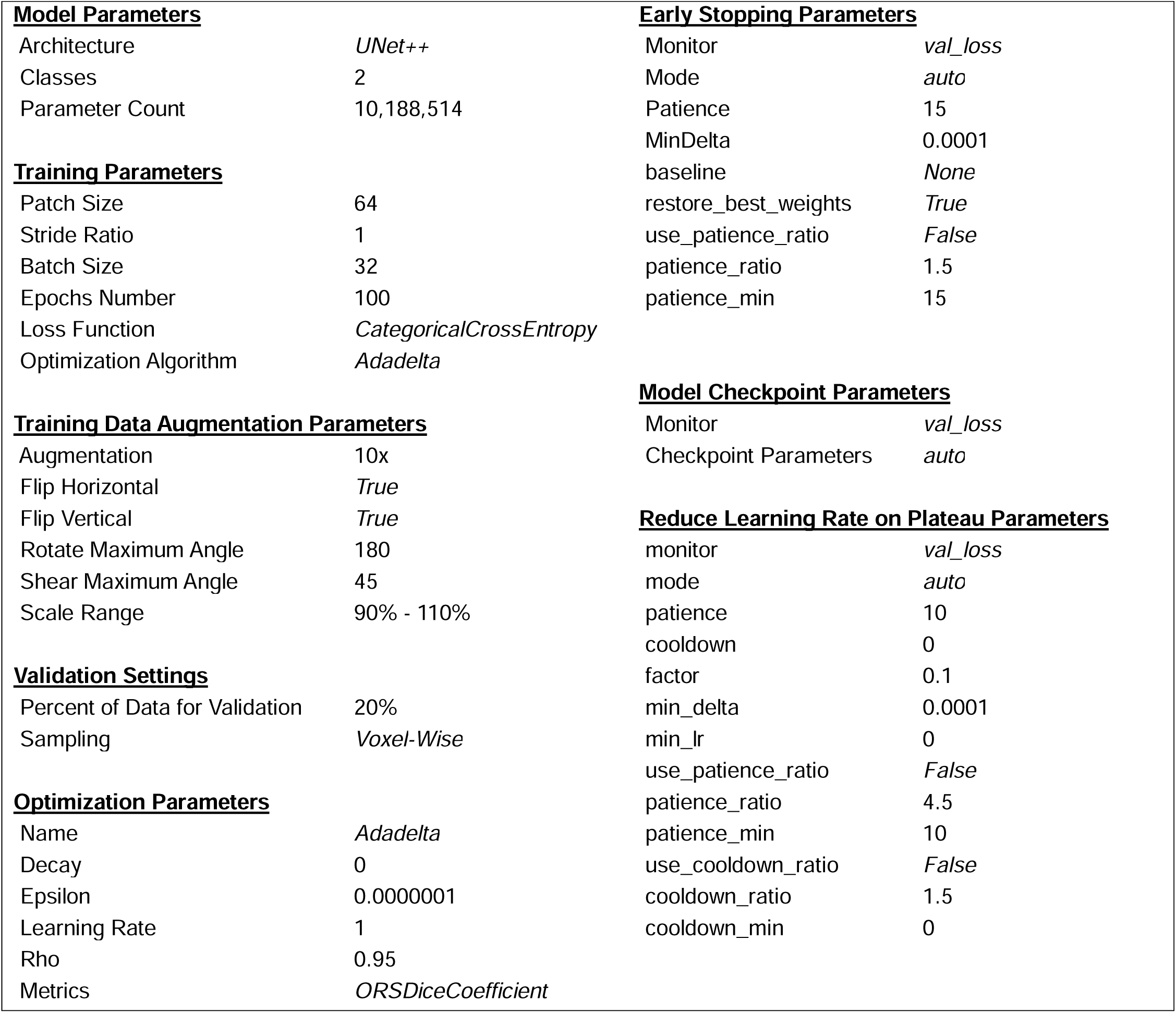
Nested UNet (UNet++) Training and Augmentation Parameters used.

